# Non-canonical proteoforms of the human fusiform gyrus in Alzheimer’s disease

**DOI:** 10.64898/2026.10.20.760462

**Authors:** Kajal Panchal, Sharmistha Majumdar

## Abstract

For a long time, the protein coding potential of our genome was underestimated. Only 2% of the genome was thought to be coding for the protein while the rest was called non-coding and junk. The two main sources of biasness which resulted in underestimation were prioritising conservation and length of the open reading frames (ORFs). But recent Ribo-seq studies and high throughput quantitative proteomics studies have shown that many ORFs previously annotated as non-coding are actually getting translated into proteins and these non-canonical proteins do have certain biological functions. These findings have added more parameter perspectives to look from about different diseases. This study aims to understand translation dynamics of these proteins encoded from non-canonical ORFs in neurological disorders affecting cognitive functions specifically targeting fusiform gyrus which is linked with face recognition and anime associations which are crucial symptoms in neurological disorders. This study uses transcriptomics and proteomics analysis to understand transcriptional and translational dynamics of ncORFs. A patient cohort from the Banner Sun Health Research Institute cohort is used, corresponding RNA-seq data and Quantitative MS/MS proteomics data published in different studies are used. A custom-made protein search database was created using transcriptomics tools like STAR, StringTie and Salmon to filter out expressed ncORFs within the non-coding region and not annotated as protein coding in either GENCODE or UniProt annotations. The custom ncORF-database was then later used in proteomics analysis as search space along with the canonical proteins. The entire comprehensive analysis has resulted in identification of 22,135 ncORF-encode proteins (ncPs) and out of which 67 are differentially expressed across different neurological conditions based on the | log2FC | >1 & p.adj < 0.05 criteria. The study highlights new molecular players which might be the missing links of known mechanisms.

## Introduction

Francois Jacob wrote in *“Evolution and Tinkering”* (1977) that, *“The probability that a functional protein would appear de novo by random association of amino acids is practically zero”*. The popular hypothesis back then was that all the genes emerge by tinkering of the previously existing ones and genomes don’t innovate new functional proteins from scratch. This hypothesis prevailed for a long time despite an early study by (Basrai et al., 1997) shed light on small open reading frames calling them beautiful needles in the haystack. Evolution by tinkering is true for many conserved protein-coding genes but ruling out any possibilities for genomic innovations kept us ignorant about the non-coding genome’s repertoire. Slowly the scepticism around these cryptic proteins got replaced by evidence driven curiosity, one of the pioneering research by (Fermin et al., 2006) presented an evidence of de novo gene emergence out of non-coding DNA in *Drosophila melanogaster* (predominantly expressed in testes, four are X-linked). Later Many species-specific de novo genes got identified, across the kingdom of life, including bacteria, plants, mice, and humans, having no ancestral link to previously known coding genes (Carvunis et al., 2012; Van Oss & Carvunis, 2019). Another dominated arbitrary assumption biased the estimation of proteome complexity, that ORFs composed of fewer than 100 amino acids are unlikely to be translated, properly folded and functional (Vrbnjak & Sewduth, 2024). Many of these de novo genes are cooked in the nursery of non-coding regions of genomes and many of them had remained unexplored which are now being explored as non-canonical open reading frames (ncORFs) translation events in many organisms, some of them being specific to lineage or tissue, developmental stage or disease conditions. High throughput technologies like Ribosome profiling and Ribo-Seq redefined analysis depths of translation further (Ingolia et al., 2009, 2011), and then the advancement in quantitative global proteomics technologies boosted the statistical confidence by validation ncORF translation events at protein levels. These ncORFs are widely known as small ORFs (smORFs) or Alternative ORFs (AltORFs), and the proteins they code are called microproteins, micropeptidespeptideins. This paper will refer to them as non-canonical ORFs (ncORFS) and ncORF-encoded proteins (ncPs) respectively. Many tissue-specific ncPs are found using proteogenomics approaches in recent studies like in mammalian heart and kidney (Flower et al., 2020; Ruiz-Orera et al., 2024; Sandmann et al., 2023), disease specific microproteins like in cancer (Callaway, 2025; Chen et al., 2025; Tornesello et al., 2025) and in immunology (Subbannayya et al., 2022). The current direction of the field is multifaceted; one is to standardize the nomenclature, cataloguing and integrating with existing annotations, towards which TransCODE Consortium is working in a large collaborative efforts (Mudge et al., 2022) and have identified 7,264 ncORF translated across different human cell types with consensus (Deutsch et al., 2026). Other aspects include characterization according to different contexts like cell type, evolutionary aspects, functionality and disease mechanisms. Most of them are evolutionarily young, emerged in the primate lineage, and are hypothesized to be responsible for many human-specific functionalities (Deutsch et al., 2024). In this study we have addressed the role of ncPs in the context of neurological disorders. The enlargement of the neocortex is thought to be responsible for advanced human cognition. (Li et al., 2023). 74 human/humanoid-specific lncRNA-originated de novo genes were identified as having a role in brain size increase over evolution (An et al., 2023; Qi et al., 2023; Rich & Carvunis, 2023; Wu et al., 2011). Ribo-seq of the developing human brain identified hundreds of previously unannotated microproteins, many from long non-coding RNAs and uORFs. A very recent study has built an atlas of ncPs of the human frontal cortex in Alzheimer’s disease (Miller et al., 2026) and have identified over 1,067 ncPs expressed in the human cortex. Few of the lncRNA encoded ncPs have been experimentally validated to have functionalities in chromatin accessibility regulation of vertebrate specific neural cells (Tornini et al., 2023) and glioblastoma cell proliferation (Du et al., 2024). Hence our study expands the understanding of neurological diseases mechanism, where we have found 67 promising ncPs which are dysregulated in the fusiform gyrus brain region of the patients with cognitive impairments. The fusiform gyrus is an important brain when it comes to measuring cognition impairment as this region handles recognition, visual processing and memory and hence the outcome of this study has relevance in diagnosis and cure.

## Methods

### Data Source

The study uses previously published datasets for transcriptomics (Friedman et al., 2018) and proteomics (Shrestha et al., 2026) analysis. Details are given in Table1.

**Table 1.** Dataset information.

| Dataset Accession ID | Publication | Description | Metadata |
| --- | --- | --- | --- |
| GEO: GSE95587 | Diverse Brain Myeloid Expression Profiles Reveal Distinct Microglial Activation | Banner Sun Health Research Institute cohort: RNA from fusiform gyrus of | Brain region: fusiform gyrus, total samples: 117,33 control, 84 Alzheimer's disease, |
|  | <u>States and Aspects of Alzheimer's Disease Not Evident in Mouse Models</u> | Alzheimer's or Neurologically Normal post-mortem tissue. | 52 females, 65 male |
| MassIVE:<br>MSV000097496<br>(AD_Cohort_1) | <u>Pan-neurodegeneration proteomics reveals disease subtypes and molecular signatures</u> | Banner Sun Health Research Institute: Global quantitative proteomics using LC-MS/MS. TMT-16 plex | Brain region: fusiform gyrus, total samples:443, Ctl:46; AD:279; MCI:77; PD:26; PSP:15 (AD: Alzheimer's disease; MCI: mild cognitive impairment; PD: Parkinson's disease; PSP: progressive supranuclear palsy; Ctl: control without dementia), Male 241, female 202 |

**Table 1.** ncORF classification criteria.

| Type | Group | Meaning |
| --- | --- | --- |
| uORF | A | Entirely in the 5' UTR |
| uoORF | A | Starts in the 5' UTR, overlaps the CDS start out of frame |
| intORF | A | Inside the CDS, different frame |
| doORF | A | Starts in the CDS, ends in the 3' UTR, different frame |
| dORF | A | Entirely in the 3' UTR |
| N_extension | A | In frame with the CDS, upstream start, same stop |
| canonical_CDS | A | The annotated CDS |
| lncRNA-ORF, retained_intron-ORF, NMD-ORF, processed_transcript-ORF, pseudogene-ORF | B | ORF in a known non-coding transcript of that type |
| intergenic-ORF | C (u) | Novel intergenic transcript |
| intronic-ORF | C (i, y) | Novel transcript inside a gene's intron |
| antisense-ORF | C (x) | Novel transcript overlapping a gene on the opposite strand |
| isoform-ORF | C (j, k, c, o) | Novel isoform of a known gene |
| retained_intron-ORF | C (m, n) | Novel model retaining an intron |
| pre-mRNA-ORF | C (e) | Single exon overlapping an intron |

### Transcriptomics Data Analysis

RNA-seq read files were downloaded from European Nucleotide Archive (ENA), and were analysed for Quality check using FastQC. Reads were aligned to reference genome GRCh38 primary assembly V50 downloaded from GENCODE (Frankish et al., 2021) using splice aware aligner STAR (Dobin et al., 2013). StringTie (M. Pertea et al., 2015) was used to assemble reads into transcript models, which output GTF file or transtomes per sample, which were then merged into a single GTF file containing all the transcript models for the entire cohort. The merged GTF file was then used to generate the Global Transcriptome FASTA file using GffRead. The same merged GTF file was used to compare assembled transcriptomes with GENCODE v50 annotations using GffCompare(G. Pertea & Pertea, 2020) package which annotates every transcript model with class code representing the transcript type compared to the reference genome annotation. These transcripts classes were used to assign ncORF types; details about the transcript class code can be found in (G. Pertea & Pertea, 2020). The derived transcripts sequence model was then used to quantify transcripts using Salmon(Patro et al., 2017) which gave output of transcript expression in TPMs. Deseq2 (Love et al., 2014) was used to analyse differential transcript expression.

### Making the search database

The main utility of transcriptomics analysis is to filter out transcripts with significant expression (>1TPM). A custom made python script was utilized to extract ncORFs hosted within novel transcript models (all GffCompare classes except s and p), known lncRNAs and other non-coding GENCODE transcripts, the UTRs of protein-coding transcripts (uORF, uoORF, dORF, doORF), out-of-frame ORFs inside coding sequences (intORF), isoform-specific ORFs, and intronic ORFs. The python script uses the filtered transcripts ids, output GTF from GffCompare which contains the comparison of built global transcripts from the cohort analysed and the reference GENECODE v50 annotation, GRch38 GENCODE 38 reference genome fasta, and reference transcriptome fasta, TPM matrix output from Salmon and canonical proteome FASTA downloaded from UniProt (2026-09-27-reviewed-isoforms-UP000005640). The ncORF extraction script takes the filtered TPM matrix and extract all the transcript ids, based on the ids it divides them into three groups A) known protein coding sequences (class code: =/eq) B) known non-coding (class code: =/eq) C) novel transcript models (class code: j,k,m,n,e,i,u,o,y,x,c) and then ORFs are extracted(from start codon AUG to stop codons), translate into amino acid sequences. Then these sequences are matched with the pool of canonical proteome to eliminate exact matched and N-terminal truncation products. The rest are kept as ncORFs and will be considered for the proteomics search database. The ncORF protein products are given comprehensive information heavy ids and the nomenclature fields are described in the Table 2. The RNA-Seq data we have used is generated from unstranded RNA-seq library preparation and hence the nucleotide sequence to amino acid translation is different for each ORFs. Only known transcripts and spliced transcripts models have reliable strand information. The script translates spliced models (with multiple exons) on their sense strand only in 3-frame translation fashion and translates single-exon models of unknown strand on both strands in 6-frame translation fashion.

**Table 2.** Putative ncORF-encoded protein ID description.

| Field | Meaning |
| --- | --- |
| ID | <transcript>_ORF<n>; ORFs numbered by position along the transcript, _rcORF<n> for the reverse-strand scan of . transcripts |
| type | ORF type (table above) |
| class | GffCompare class code (eq ==) |
| gene, gene_id | Gene name and ID (reference gene when known, else the StringTie gene) |
| tx, ref_tx, ref_tx_type | Transcript scanned, matched reference transcript, its GENCODE type |
| loc | chr:start-end:strand, 1-based genomic span of the ORF including the stop codon |
| blocks | Number of exon pieces the ORF spans |
| tx_exons | Exons in the transcript (1 = single-exon model) |
| tx_pos, tx_len | ORF start–end within the transcript (1-based, stop included) and transcript length |
| tx_strand, scan | Transcript strand (+, -, .) and strand read (+ sense, rc reverse complement) |
| len_aa, start | Protein length without stop; start codon used |
| maxTPM | Highest TPM of the transcript across samples |
| match | Comparison with the canonical proteome (novel or canonical_N_extension) |
| n_sources | ORF copies with this identical peptide |

### Proteomics Analysis

Thermo Fisher raw files were downloaded from the MassIVE database. ThermoRawFileParser(Hulstaert et al., 2020) was used to convert raw files into mzml file format. For proteomics analysis we used FragPipe v24.0 (*GitHub - Nesvilab/FragPipe*, n.d.), the workflow was chosen as TMT-16 as the proteomics data was originally proposed by TMT-16 plex method. Most of the default parameters were used, Trypsin was selected as the digestion enzyme as per the experiment. Methionine oxidation and protein N-terminal acetyl variable modifications and carbamidomethyl-fixed modifications were used.peptide length (aa) range was kept from 4 to 100 and peptide mass range (Da) was kept from 200 - 5000. The custom made ncORF protein search database was used along with the canonical UniProt reviewed proteome dataset with isoforms UP000005640 and common protein contamination. In the final search database, the canonical proteome consists of about 42,547 protein sequences while putative ncORF-encoded proteins (ncPs) were 10,99,695. After addition of common contaminant proteins and decoys (50%) total protein sequences were 21,99,486. Decoy sequences are simply the reverse sequences of the database proteins which will be used to calculate the false discovery rate (FDR) of the database hits, to filter out statistically significant positives. FragPipe is a proteomics analysis pipeline, which includes MsFragger (Kong et al., 2017) search engine that matches experimental spectra with that of the theoretical spectra and identifies spectra-peptide match, Philosopher toolkit (da Veiga Leprevost et al., 2020) performs the downstream analysis of the MsFragger matched to filter out based on FDR (retailing matches with <0.0.1 FDR), IonQuant (Yu et al., 2021) which quantifies the protein expression with help of TMT-integration (Chang et al., 2026)for TMT workflows. The result of the FragPipe is a quantification report containing proteins identified.

### Differential Protein Expression Analysis

FragPipeAnlaystR (Hsiao et al., 2024) an R package was used for the differentially expressed protein expression analysis which uses limma. Significance cut off was taken as | log2FC | > 1 and, p.adj < 0.05. Input files were ratio_protein_MD.tsv output from tmt-integrator which contains Index NumberPSM, Protein, MaxPepProb, Protein Protein ID, Entry Name, Protein Description, Organism, Indistinguishable, Protein, ReferenceIntensity and experiment_annotation.tsv file which contains metadata of the samples and which TMT channel corresponds to which sample and experiment condition.

## Results

### Non-canonical ORF-encode proteins (ncPs) are found to be dysregulated in neurological disease patients postmortem fusiform gyrus

A total of 21,9104 proteins were detected by fragpipe (expressed at protein level), out of which 22,135 are novel ncORF encoded proteins.Total of 468 differentially expressed (DE) proteins were found using the statistical method called limma and out of that 67 were DE ncPs across 7 different condition comparisons, both up and down regulations has been observed. There were 5 different conditions; Alzheimer’s disease (AD); mild cognitive impairment (MCI), Parkinson’s disease (PD), progressive supranuclear palsy (PSP),control without dementia (Ctl). Variety of ORF types have been identified to be expressed and differentially expressed, among which novel isoform ORFs are dominant. Figure 2 represents the number distribution across the ncORF types and comparison types. Figure 3 shows the different features of DE ncPs, like their host transcript max expression across samples (max TP), LogFC values distribution, max PepProb, Reference Intensity and chromosome distribution.

**Figure 1.**
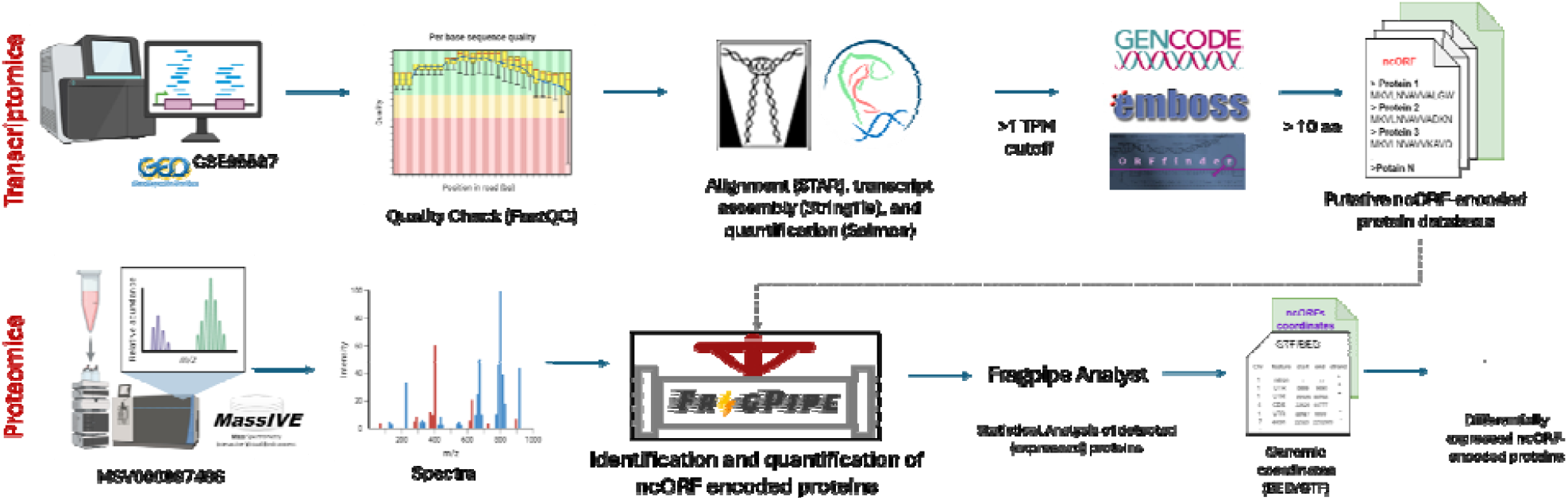
Overall Workflow

**Figure 2.**
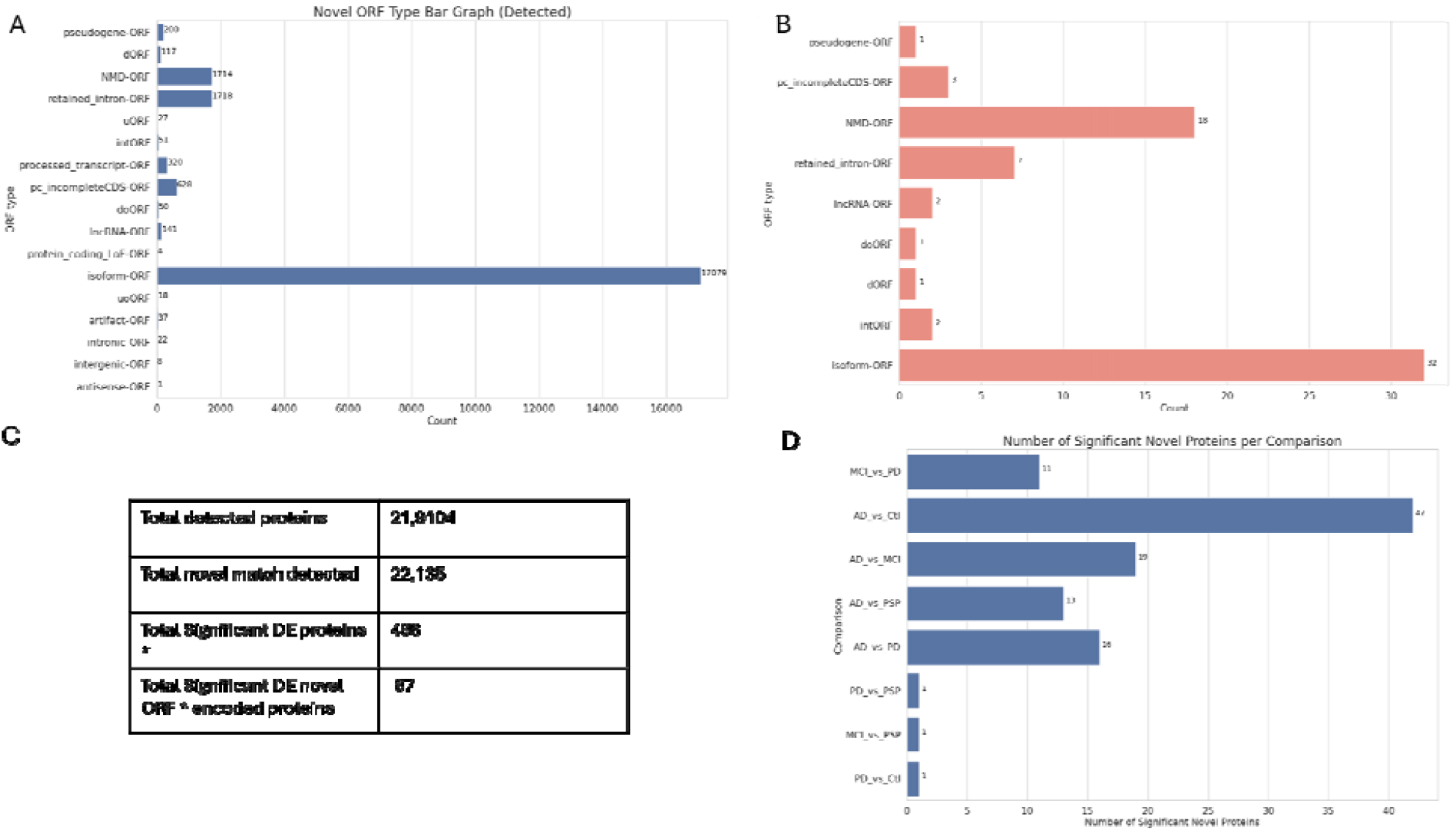
A) ncORF type distribution of the novel ncORF-encoded proteins expressed pool B) ncORF type distribution of the novel ncORF-encoded proteins differentially expressed pool (DE) C)Numbers of proteins detected and differentially expressed based on Fragpipe analysis (criteria: *| log2FC | >, p.adj < 0.05) D) DE-ncPs across differentially condition comparisons

**Figure 3.**
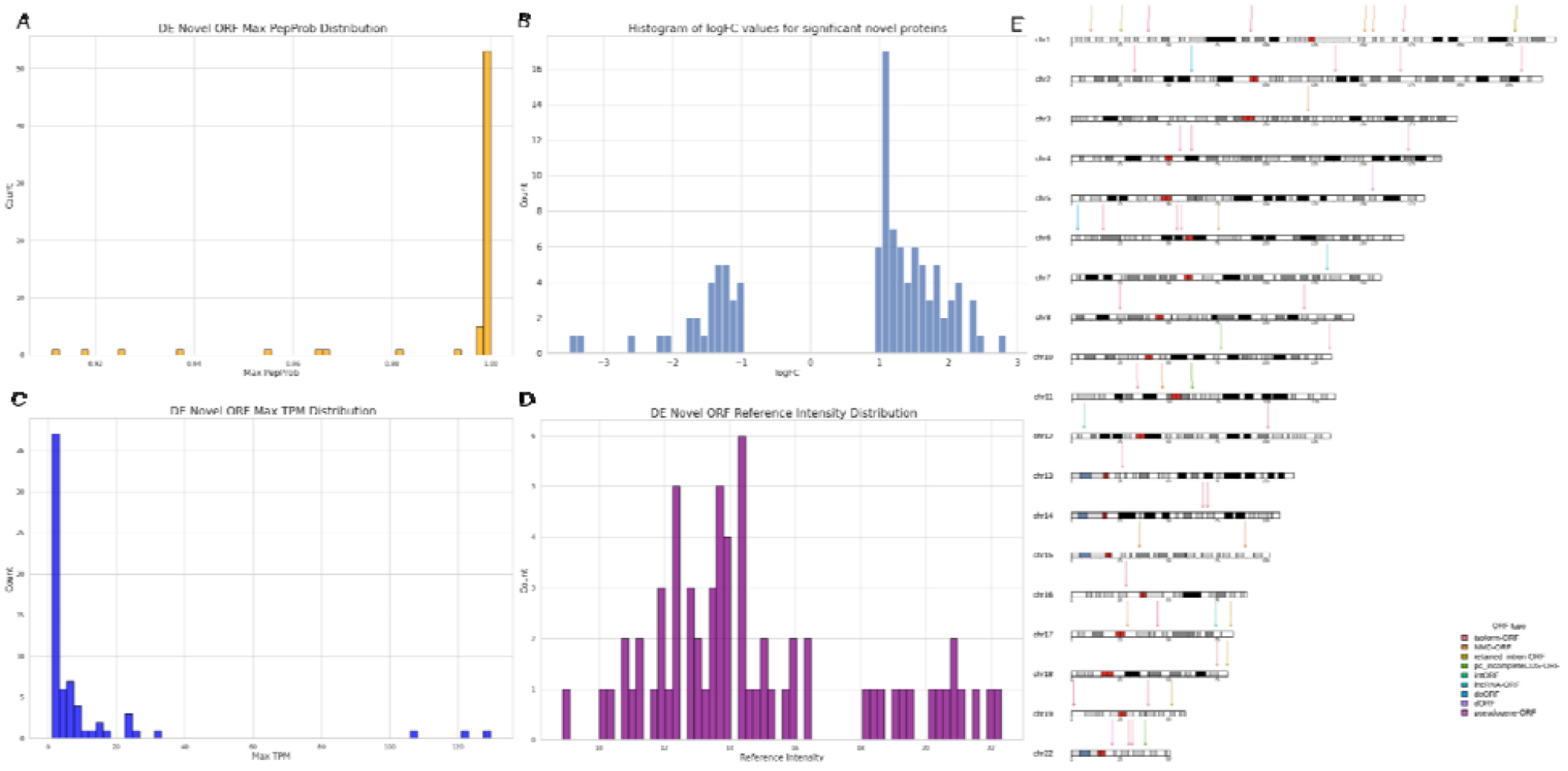
A) DE novel ncP max PepProb distribution B) log2FC value distribution of DE novel ncORF-encoded proteins C) DE novel ncPs max TPM distribution D) DE ncP reference intensity distribution E) DE ncPs chromosomal distribution visualization across different types of ncORFs

### Dysregulated ncPs’ host transcripts are not found to be dysregulated at transcript level

Differential expression analysis of transcripts were also performed, although there was no overlap between DE novel transcripts and host transcripts of DE-ncPs.

## Conclusions

Novel ncORF encoded proteins are indeed getting translated as well as dysregulated and thus might have a role to play in neurological disorders and could become potential targets for diagnosis and therapeutics.

